# Siglec-1 on dendritic cells mediates SARS-CoV-2 *trans*-infection of target cells while on macrophages triggers proinflammatory responses

**DOI:** 10.1101/2021.05.11.443572

**Authors:** Daniel Perez-Zsolt, Jordana Muñoz-Basagoiti, Jordi Rodon, Marc Elousa, Dàlia Raïch-Regué, Cristina Risco, Martin Sachse, Maria Pino, Sanjeev Gumber, Mirko Paiardini, Jakub Chojnacki, Itziar Erkizia, Xabier Muñiz, Ester Ballana, Eva Riveira-Muñoz, Marc Noguera, Roger Paredes, Benjamin Trinité, Ferran Tarrés-Freixas, Ignacio Blanco, Victor Guallar, Jorge Carrillo, Julià Blanco, Amalio Telenti, Holger Heyn, Joaquim Segalés, Bonaventura Clotet, Javier Martinez-Picado, Júlia Vergara-Alert, Nuria Izquierdo-Useros

**Author notes:** Equal contributions (Daniel Perez-Zsolt, Jordana Muñoz-Basagoiti, Jordi Rodon, Marc Elousa). senior authorships (Javier Martinez-Picado, Júlia Vergara-Alert, Nuria Izquierdo-Useros). **Corresponding authors**: JM-P; JV-A and NI-U.

## Abstract

COVID-19 pandemic is not yet under control by vaccination, and effective antivirals are critical for preparedness. Here we report that macrophages and dendritic cells, key antigen presenting myeloid cells (APCs), are largely resistant to SARS-CoV-2 infection. APCs effectively captured viruses within cellular compartments that lead to antigen degradation. Macrophages sense SARS-CoV-2 and released higher levels of cytokines, including those related to cytokine storm in severe COVID-19. The sialic acid-binding Ig-like lectin 1 (Siglec-1/CD169) present on APCs, which interacts with sialylated gangliosides on membranes of retroviruses or filoviruses, also binds SARS-CoV-2 via GM1. Blockage of Siglec-1 receptors by monoclonal antibodies reduces SARS-CoV-2 uptake and transfer to susceptible target cells. APCs expressing Siglec-1 and carrying SARS-CoV-2 are found in pulmonary tissues of non-human primates. Single cell analysis reveals the *in vivo* induction of cytokines in those macrophages. Targeting Siglec-1 could offer cross-protection against SARS-CoV-2 and other enveloped viruses that exploit APCs for viral dissemination, including those yet to come in future outbreaks.

## INTRODUCTION

Despite the efficacy and progress of vaccines against severe acute respiratory syndrome coronavirus 2 (SARS-CoV-2), future viral variants and other coronaviruses with similar outbreak potential might overcome these strategies and will require treatments yet to be developed. Since global immunization campaigns will likely require years to cover all the population worldwide, there is an urgent need to implement effective therapeutic treatments to reduce the mortality associated with severe coronavirus infectious disease 2019 (COVID-19). The respiratory illness caused by SARS-CoV-2 leads to hospitalization in 10-30% of the infected individuals and, eventually, the admission into the intensive care unit (CDC Weekly, 2020; European Center for Disease Prevention and Control, 2020). Severe COVID-19 is associated with pneumonia, dyspnea, hypoxemia and lymphopenia, and can rapidly progress to respiratory failure (Berlin et al., 2020). Acute respiratory distress syndrome (ARDS) is also a common complication and is associated with a disproportionate inflammatory response to SARS-CoV-2, characterized by a cytokine storm-like syndrome (Hadjadj et al., 2020; Huang et al., 2020; Mehta et al., 2020; Valle, 2020). Of note, mortality rate in COVID-19-associated ARDS is 45%, and the incidence of ARDS among non-survivors of COVID-19 is 90% (Tzotzos et al., 2020).

Earlier studies have highlighted the paramount role of myeloid antigen-presenting cells (APCs), such as macrophages and dendritic cells (DCs), in mediating an antiviral inflammatory response, which is exacerbated in severe COVID-19 cases (Bain et al., 2021; Merad and Martin, 2020). A critical role of lung macrophages in inducing the inflammation associated with the pathologic sequelae of SARS-CoV-2 infection has been confirmed also in nonhuman primates (NHP) (Hoang et al., 2021). Yet, whether these cells effectively trap and process SARS-CoV-2, and if their viral detection results in the secretion of inflammatory cytokines need further investigation. Myeloid APCs might contribute to viral pathogenesis if they are susceptible to infection, or able to capture and transfer infectious viruses to bystander target cells, thereby favoring viral spread to other tissues (Geijtenbeek et al., 2000; Moller-Tank and Maury, 2015; Sewald et al., 2016). Indeed, viral dissemination mediated by APCs is a common pathway co-opted by different types of viruses to evade immunity, as it is the case of the human immunodeficiency virus type 1 (HIV-1), which is effectively transferred to target cells via a mechanism known as *trans*-infection (Cameron et al., 1992; Geijtenbeek et al., 2000). For other viruses, such as Ebola virus, productive infection of APCs is critical in determining host susceptibility and viral dissemination to distant tissues (Geisbert et al., 2003; Martines et al., 2015).

Early interactions between viruses and macrophages or DCs are critical to either trigger immune responses or favor viral dissemination. For SARS-CoV-2, the outcome of such interactions has not been fully explored, although it might impact the progression of the COVID-19 severity. Thus, elucidating the initial steps of viral interaction with myeloid APCs is essential for the design of new therapies to increase protection of exposed individuals. Several lectin receptors are critical for initial viral recognition on APCs. C-type lectins such as DC-specific intercellular adhesion molecule-3-Grabbing non-integrin (DC-SIGN) mediate the attachment of several viruses such as HIV-1 or Ebola virus via viral glycoprotein recognition (Alvarez et al., 2002; Geijtenbeek et al., 2000; Lin et al., 2003; Simmons et al., 2003), being also the case for SARS-CoV-2 (Thépaut et al., 2020). The sialic acid-binding lectin Siglec-1 is an interferon inducible receptor expressed on activated myeloid cells (Hartnell et al., 2001; Pino et al., 2015; Puryear et al., 2013) whose expression is up-regulated on APCs in SARS-CoV-2 infected individuals (Bedin et al., 2020). Siglec-1 is implicated in the binding of HIV-1 and Ebola virus via the recognition of sialylated ligands exposed on the lipid membranes of these viruses. The V-set domain of Siglec-1 interacts with sialyllactose on viral membrane gangliosides (Izquierdo-Useros et al., 2012a, 2012b; Puryear et al., 2012, 2013), which HIV-1, Ebola virus and other enveloped viruses incorporate during the budding from the membranes of infected cells (Bavari et al., 2002; Chan et al., 2008; Feizpour et al., 2015; Izquierdo-Useros et al., 2012a; Kalvodova et al., 2009). However, a general role for Siglec-1 in facilitating SARS-CoV-2 uptake and *trans*-infection to pulmonary target cells such as pneumocytes or respiratory ciliated cells remains largely unexplored, despite being one of the lectins that are highly expressed on pulmonary macrophages.

Here we show that, although being largely resistant to SARS-CoV-2 infection, myeloid cells effectively capture and trap incoming viruses in internal compartments connected with the plasma membrane, eventually leading to viral degradation overtime. Among the myeloid cells tested, macrophages released higher amounts of inflammatory cytokines upon viral sensing, including those involved in the cytokine storm associated to ARDS (Hadjadj et al., 2020; Huang et al., 2020; Mehta et al., 2020; Valle, 2020). Finally, we show that Siglec-1 mediates SARS-CoV-2 recognition of different viral variants of concern via interaction with sialylated ligands, such as the ganglioside GM1 identified on SARS-CoV-2 membrane. Immunohistochemistry and single cell RNA analyses of pulmonary tissues of SARS-CoV-2 infected NHP corroborated, directly *in vivo*, the presence of Siglec-1 on myeloid cells containing viruses. Siglec-1 capacity to bind SARS-CoV-2 was more relevant than other well-known attachment receptors such as DC-SIGN, and more effective at mediating transfer of viruses to susceptible target cells via *trans*-infection. Since anti-Siglec-1 monoclonal antibodies (mAbs) blocked SARS-CoV-2 *trans*-infection, targeting Siglec-1 could offer cross-protection against different SARS-CoV-2 variants and other enveloped viruses that exploit APCs for viral dissemination.

## RESULTS

### 1. Myeloid cells are not productively infected by SARS-CoV-2 but capture and degrade trapped viruses

Preliminary reports indicate that myeloid cells such as monocyte-derived macrophages and DCs are highly resistant to coronavirus infection (Dalskov et al., 2020; Tynell et al., 2016; Yilla et al., 2005). In line with these findings, we observed a reduction above 50% in the viability of susceptible Vero E6 cells 3 days post SARS-CoV-2 infection, that was not observed for monocyte-derived myeloid cells (**Figure 1A**). Of note, upon SARS-CoV-2 invasion, myeloid cells responded to activating signals that initiate antiviral responses, such as the release of type I interferon-α (IFN-α), or to the presence of bacterial lipopolysaccharide (LPS), which are increased throughout the course of SARS-CoV-2 infection (Giron, 2021). Thus, we tested the fusion capacity of SARS-CoV-2 spike on myeloid cells activated or not in the presence of IFN-α. Reporter pseudoviral vectors expressing VSV glycoprotein effectively fused with both macrophages and DCs, although it decreased on IFN-α treated myeloid cells (**Figure 1B**). However, fusion was always negligeable when pseudoviruses were pseudotyped with SARS-CoV-2 spike. Of note, both pseudoviruses effectively fused with HEK-293T cells expressing ACE2. For SARS-CoV-2 spike pseudovirus this was specifically blocked by a human ACE2-Fc fusion protein, which was not active on myeloid cells (**Figure 1B**).

**Figure 1.**
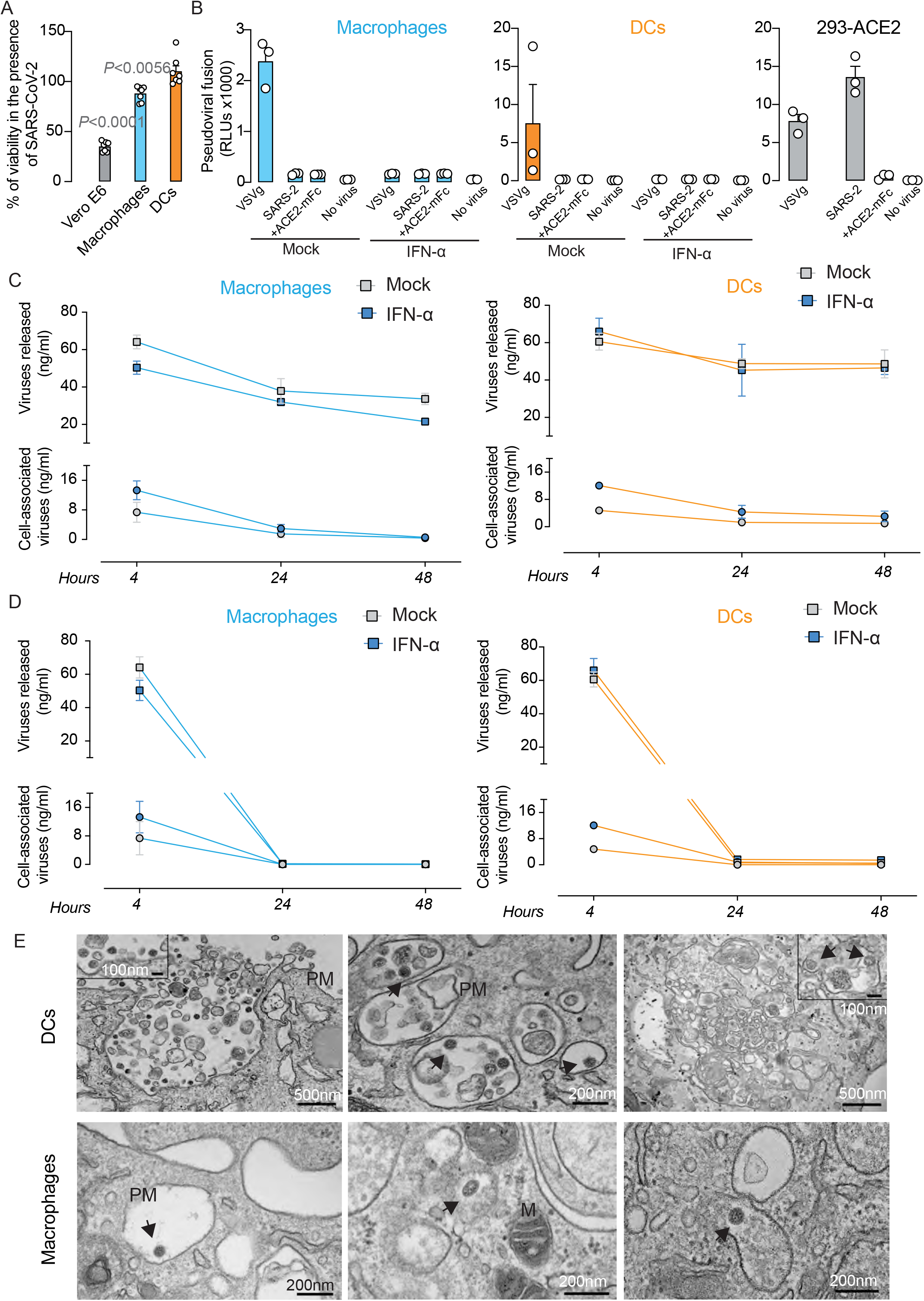
Myeloid cells are not productively infected by SARS-CoV-2 but these APCs capture and degrade trapped viruses. **A**. Percentage of cellular viability 3 days post infection with SARS-CoV-2 at a MOI of 0.1 in Vero E6 cells, macrophages and DCs. Values from 2 replicates and four experiments. Statistical differences from 100% of viability were assessed with a one-sample t-test. **B**. Fusion of HIV-1 luciferase reporter viruses lacking the envelope glycoprotein pseudotyped with VSV glycoprotein or SARS-CoV-2 Spike in macrophages and DCs stimulated or not with IFN-α and in ACE2 expressing HEK-293T cells. ACE2-mFc fusion protein was used to block ACE2-dependent viral fusion. Values from 2 replicates and one experiment. **C**. Uptake of SARS-CoV-2 by macrophages (left graph) and DCs (right graph) activated or not with IFN-α that were pulsed for 4 h, 24 h and 48 h at 37 °C with the virus, to assess the amount of virus present in the supernatant (squares) and the amount of cell-associated viral nucleocapsid detected on cellular lysates after extensive washing (circles) by ELISA. Data from one representative experiment out of two shows means and SEM and from 3 different donors. **D**. Kinetics of SARS-CoV-2 degradation after 4 h of viral exposure and extensive washing by macrophages (left graph) and DCs (right graph) activated or not with IFN-α. The amount of virus present in the supernatant (squares) and the amount of cell-associated viral nucleocapsid detected on cellular lysates after extensive washing (circles) was measured by ELISA. Data from one representative experiment out of two shows means and SEM from 3 different donors. **E**. Electron microscopy images of Macrophages and LPS DCs exposed first to SARS-CoV-2 at an MOI of 1. Arrows indicate individual viral particles. PM, plasma membrane, M, mitochondrion. Data from 2 donors and 2 experiments.

Next, we compared the capacity of myeloid cells activated or not in the presence of IFN-α to capture SARS-CoV-2 by an ELISA measuring the amount of cell-associated virus. We followed the fate of these viruses for two days, assessing the amount of cell-associated virus captured overtime while measuring the amount of virus remaining in the supernatant (**Figure 1C**). Maximal viral uptake on cells was detected 4 h post viral addition, and decreased overtime for activated and non-activated myeloid cells, indicating that APCs were effectively processing captured viruses, since SARS-CoV-2 detected in the supernatant also diminished over time (**Figure 1C**). After extensive cellular washing, degradation of viruses trapped for 4 h was also assessed on macrophages and DCs previously activated or not with IFNα (**Figure 1D**). Cell-associated viruses detected at 4 h were quickly degraded after 24 h of culture, confirming that APCs were effectively processing captured viruses (**Figure 1D**). This was not due to the release of bound viruses to the supernatant, where no SARS-CoV-2 accumulation could be detected over time (**Figure 1D**). Moreover, the absence of viral secretion to the media confirmed the inefficient viral replication of SARS-CoV-2 in myeloid cells regardless of their activation status (**Figure 1D**). In agreement with these results, electron microscopy micrographs of myeloid cells showed SARS-CoV-2 particles associated with sack-like structures resembling viral containing compartments (VCC) on APCs (**Figure 1E**), as already described for other viruses (Perez-Zsolt et al., 2019; Yu et al., 2008). On DCs (**Figure 1E**, first two images on the top) we found extracellular viruses attached to invaginations of the plasma membrane or VCCs that appear also as vacuoles. These structures were however connected to the plasma membrane. In addition, we found viruses on large membranous compartments resembling degradative structures, where damaged viral particles were observed (**Figure 1E**, last image on the top). On Macrophages (**Figure 1E**, first image on the bottom) we also found extracellular viruses attached to the plasma membrane, but also on vesicles filled with material that marks them as endocytic structures (**Figure 1E**, last two images on the bottom). Hence, viruses were effectively captured by APCs, especially upon cellular activation, that were in any case effectively processed by all APCs.

### 2. SARS-CoV-2 is effectively sensed by macrophages that release increased levels of cytokines as compared to DCs

We then sought to see if SARS-CoV-2 captured and processed by non-activated monocyte-derived myeloid cells could induce cytokine release upon initial viral sensing using Luminex. The fold change in the secretion of cytokines and chemokines by myeloid cells exposed to SARS-CoV-2 as compared to those left untreated was measured over time. One day after SARS-CoV-2 exposure, macrophages released increased levels of IL-6 (**Figure 2A**) and cytokine secretion was maintained at 48 h. A much lower viral-induced release of IL-6 was observed for DCs at 24 h and was lost overtime (**Figure 2A**). Similar dynamics were observed for tumor necrosis factor-α (TNF-α) and macrophage inflammatory protein-1α (MIP-1α) on DCs. In macrophages, TNF-α was also induced 2-fold upon SARS-CoV-2 exposure 4 h and 24 h after viral exposition, and above six-fold at 48 h (**Figure 2A**). MIP-1α was up-regulated 4 h after viral exposure in macrophages, but the release of this chemokine decreased to basal levels at 48 h (**Figure 2A**). Interferon-γ inducible protein 10 (IP-10) was induced by SARS-CoV-2 over time in macrophages, and in this particular case, the induction was higher for DCs than for macrophages (**Figure 2A**). These data indicate that macrophages sensed and responded to SARS-CoV-2 more effectively than DCs, although both cellular types efficiently engulfed and degraded these viruses.

**Figure 2.**
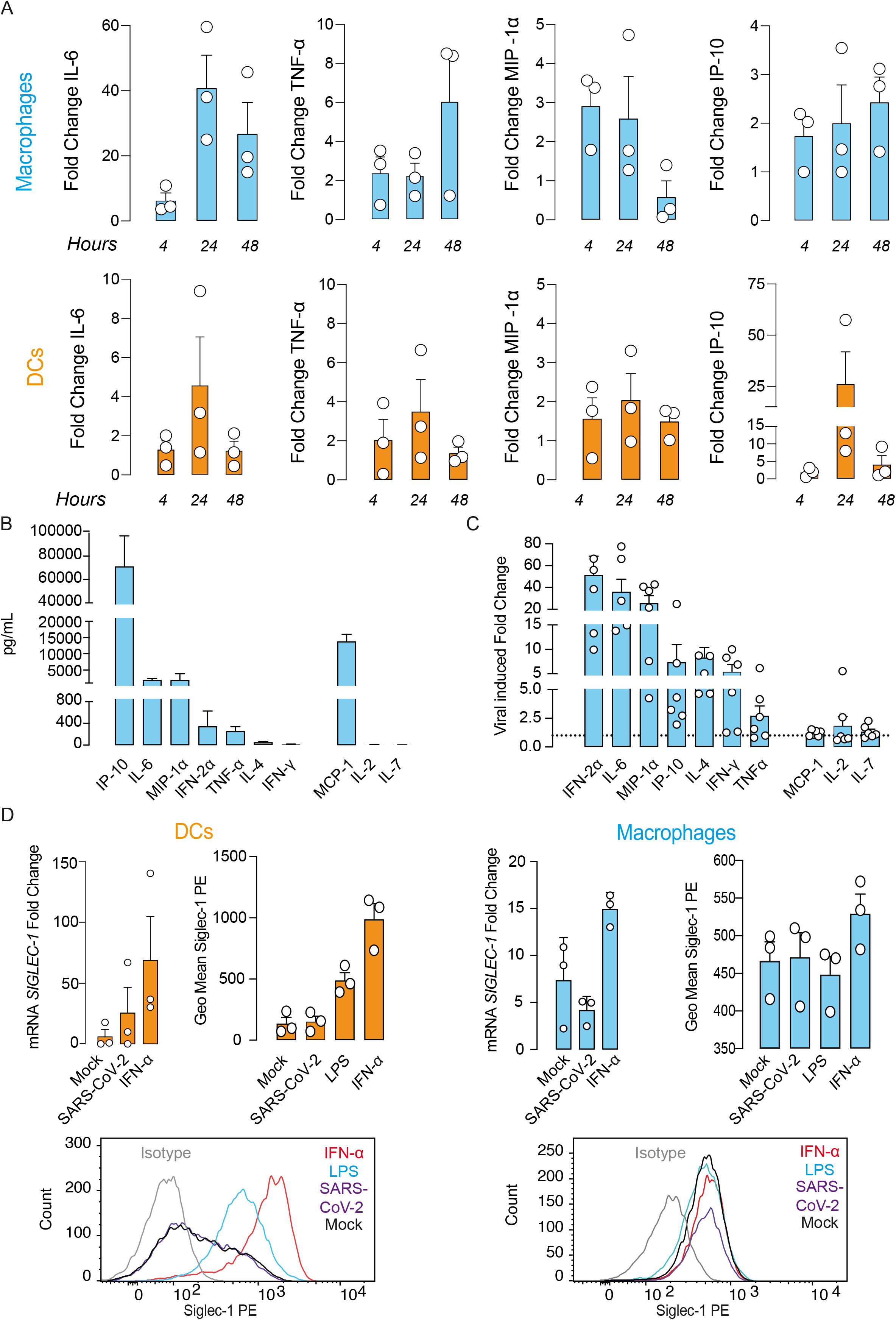
SARS-CoV-2 is effectively sensed by macrophages. **A**. Fold change in cytokine release of macrophages and DCs as compared to mock-treated cells 4 h, 24 h and 48 h post viral addition. Data from cells from 3 different donors. **B**. Quantity of cytokines released by macrophages exposed to SARS-CoV-2 for 24h at a MOI of 0.1. Data show mean values and SD of duplicate measurements of cells from 3 different donors. **C**. Fold change in cytokine release of macrophages in B, as compared to mock-treated cells. Data show duplicate measurements of cells from 3 different donors. **D**. Fold change on *SIGLEC1* mRNA induction after 24h of exposure to IFN-α or SARS-CoV-2 (MOI=0.01) and representative Siglec-1 surface staining of myeloid cells equally activated and also exposed to LPS analyzed by FACS. Results from six independent biological replicates and two experiments.

We therefore focused on macrophages, and quantified the concentration of cytokines released one day after viral exposure, and the fold change in cytokine secretion as compared to mock-treated cells (**Figure 2B**). Among the viral-induced cytokines, the most abundantly secreted proteins increased after macrophage viral sensing were IP-10, followed by IL-6, MIP-1 α, IFN-2α, TNF-α, and then by IL-4 and IFN-γ (**Figure 2B**). Of those, IFN-2α, IL-6, and MIP-1α increased more than 20 times as compared to untreated macrophages (**Figure 2C**). MCP-1, IL-2 and IL-7 secretion were however not increased by SARS-CoV-2 exposure (**Figure 2C**). Thus, among myeloid cells, macrophages secreted higher levels of cytokines upon SARS-CoV-2 recognition compared to DCs. While released interferons may have antiviral properties, cytokines such as IL-6, MIP-1α or TNF-α are involved in the disproportionate inflammatory storm-like response against SARS-CoV-2 that is associated to the ARDS complication of COVID-19 (Hadjadj et al., 2020; Huang et al., 2020; Mehta et al., 2020; Valle, 2020).

To assess if SARS-CoV-2 could induce activation of myeloid cells, we pulsed DCs and macrophages for 48 h with SARS-CoV-2, LPS or IFN-α. On DCs, LPS triggered upregulation of the human leukocyte antigen (HLA)-DR and co-stimulatory molecules, but decreased the levels of DC-SIGN (**Supplementary Figure 1**). Yet, SARS-CoV-2 exposure did not activate DCs to the same extent (**Supplementary Figure 1**). On macrophages, none of these stimuli nor the virus impacted on the expression of HLA-DR or co-stimulatory molecules (**Supplementary Figure 1**). Of note, *SIGLEC1* mRNA induction was always more potently triggered by IFN-α than by SARS-CoV-2 (**Figure 2D**), which correlated with Siglec-1 expression on the cellular surface (**Figure 2D**), both on DCs and macrophages. These results indicate that direct exposure to SARS-CoV-2 does not activate DCs as potently as LPS or IFN-α. Yet, these bystander activation stimuli released throughout SARS-CoV-2 infection can induce Siglec-1 expression on APCs, especially on DCs (**Figure 2C and D**).

### 3. Siglec-1 receptor on APCs binds SARS-CoV-2 variants via sialic acid recognition present on viral membrane gangliosides

Siglec-1 is upregulated on myeloid cells upon SARS-CoV-2 infection (Bedin et al., 2020), and strongly correlates with SARS-CoV-2 RNA counts on single cell analysis of bronchoalveolar lavages from COVID-19 patients (Lempp et al., 2021). Siglec-1 is involved in the uptake by APCs of different viruses (including HIV-1 and Ebola virus) via recognition of syalilated gangliosides anchored on the viral membranes (Bavari et al., 2002; Chan et al., 2008; Feizpour et al., 2015; Izquierdo-Useros et al., 2012a; Lorizate and Kräusslich, 2011; Panchal et al., 2003). We therefore tested if Siglec-1 could recognize and capture SARS-CoV-2 by assessing viral uptake in two complementary cellular models. First, we used the antigen presenting Raji B cell line transfected with different lectins, whose level of expression is shown in **Supplementary Figure 2**. We measured the capacity of these cells for SARS-CoV-2 uptake. All Raji cells were pulsed with equal amounts of SARS-CoV-2, extensively washed, lysed and assessed by ELISA to measure the amount of viral nucleocapsid protein. While Raji Siglec-1 cells effectively captured SARS-CoV-2, Raji cells transfected with Siglec-5, Siglec-7, DC-SIGN or devoid of any of these lectins did not (**Figure 3A**). We next tested if Siglec-1 uptake of SARS-CoV-2 relied on the recognition of sialylated ligands, which most likely are gangliosides exposed on viral membranes, as previously described for HIV-1 and Ebola virus (Izquierdo-Useros et al., 2012a, 2012b; Perez-Zsolt et al., 2019; Puryear et al., 2012, 2013). This was confirmed using a Raji cell line transfected with the Siglec-1 mutant R116A, which contains a mutation critical for sialic acid recognition (Hartnell et al., 2001; Puryear et al., 2013) that did not trap SARS-CoV-2 (**Figure 3A**), as previously shown for other viruses (Perez-Zsolt et al., 2019; Puryear et al., 2013). To further confirm if SARS-CoV-2 interaction was mediated by Siglec-1, we pre-treated Raji cells with an α-Siglec-1 mAb 7-239 previously shown to decrease HIV-1 and Ebola virus uptake (Izquierdo-Useros et al., 2012b; Perez-Zsolt et al., 2019; Puryear et al., 2013). While isotype mAb had no inhibitory effect, pre-treatment with 7-239 mAb clearly reduced SARS-CoV-2 uptake (**Figure 3B**). Of note, distinct SARS-CoV-2 variants (D614G, B.1.1.7 first identified in UK, B.1.1.248.2 first identified in Brazil, and the B.1.351 first identified in South Africa) were equally trapped via Siglec-1 receptor, but were not captured by the mutated Siglec-1 R116A, indicating that sialic acid recognition is critical for viral trapping (**Figure 3C**).

**Figure 3.**
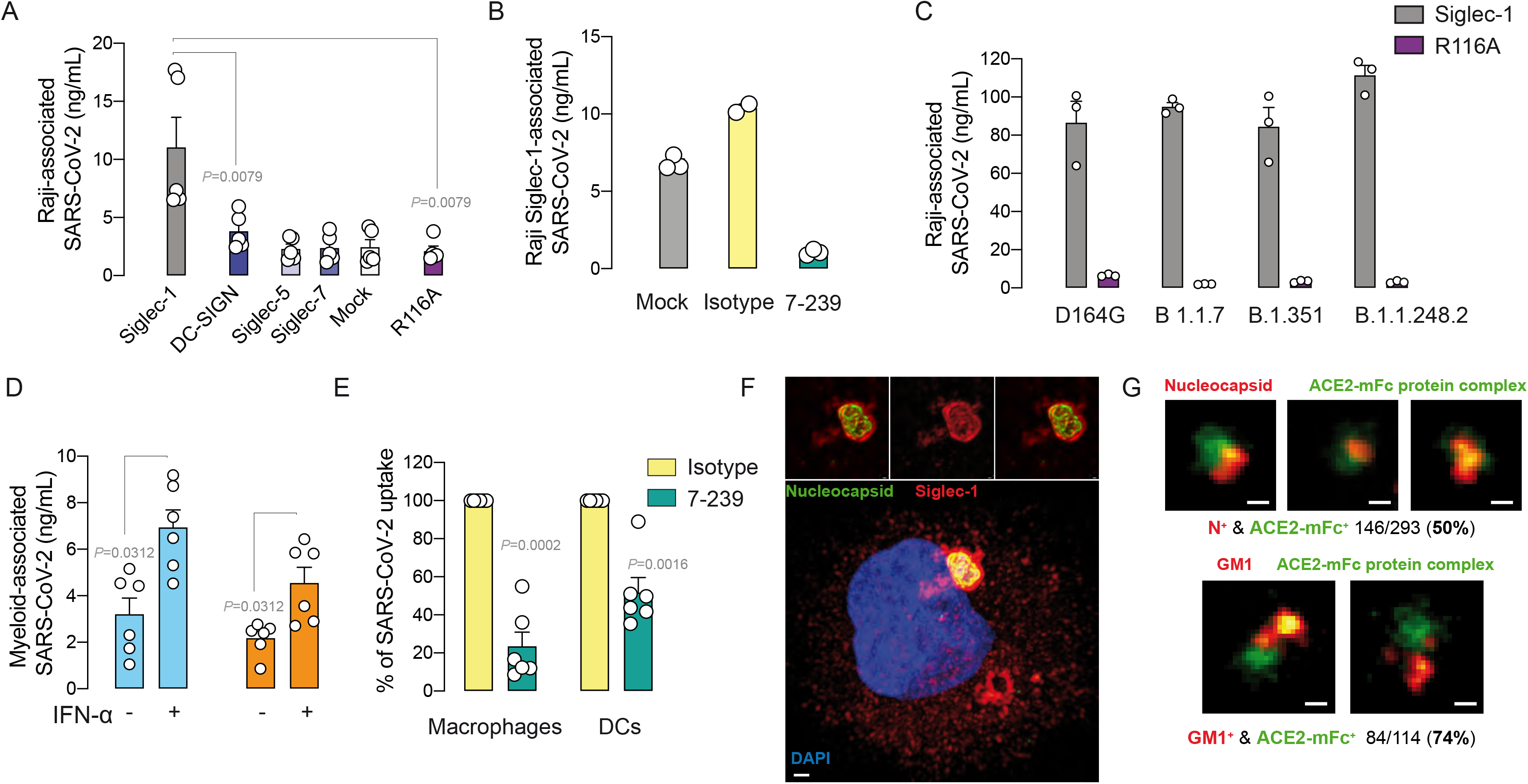
Siglec-1 receptor binds SARS-CoV-2 variants via sialic acid recognition present on viral membrane gangliosides. **A**. Comparative uptake of SARS-CoV-2 by distinct Raji B cells, that were pulsed for 2 h at 37°C, washed and lysed to assess the amount of cell-associated viral nucleocapsid by ELISA. Values from five replicates and two experiments. Statistical differences were assessed with a Mann Whitney *t* test. **B**. Uptake of SARS-CoV-2 by Raji Siglec-1 pre-incubated with α-Siglec-1 mAb 7239 or the corresponding isotype control processed as in A. Values from three replicates and one experiment. **C**. Comparative uptake of 4 different variants of SARS-CoV-2 by Raji Siglec-1 and Raji R116A Siglec-1 B cells that were processed as in A. **D**. Uptake of SARS-CoV-2 by macrophages and DCs activated or not with IFNα for 4 h and then processed as in A. Values from at least three donors and two experiment. Statistical differences were assessed with a Wilcoxon matched paired *t* test. **E.** Percentage of viral uptake inhibition of macrophages and DCs activated IFNα that were pre-incubated with 10 μg/mL of the indicated mAbs and pulsed with SARS-CoV-2 (MOI=0.1). **F**. Confocal microscopy image of an LPS-treated DC exposed to SARS-CoV-2 at a MOI of 1 showing a 3D reconstruction of the VCC and the whole cell (Scale bars 0,2 μM and 1 μM, respectively). Cells were stained with anti-nucleocapsid pAbs (green), anti-Siglec-1 mAb (red) and Dapi (blue) to stain the nucleus. See also supplementary movie 1. **G**. Super resolution microscopy of SARS-CoV-2. Top images: viruses stained with anti-nucleocapsid Abs (red) and ACE2-mFc fusion protein that interacts with the Spike of SARS-CoV-2 (green). Bottom images: viruses stained with anti-GM1 Abs (red) and ACE2-mFc fusion protein that interacts with the Spike of SARS-CoV-2 (green). Percentage of co-staining for each is shown. Scale bar: 100 nm.

We also used a second cellular model of monocyte-derived macrophages and DCs to verify the role of Siglec-1 receptor on primary cells. Myeloid cells previously treated or not with IFN-α were pulsed with SARS-CoV-2, washed and assayed by ELISA as described for Raji cells. Of note, all myeloid cells trapped SARS-CoV-2 but IFN-α activated APCs, which display higher amounts of Siglec-1 (**Figure 2D**), were the cells with higher uptake capacity (**Figure 3D**). To further investigate whether SARS-CoV-2 viral interaction was mediated by Siglec-1, we used the anti-Siglec-1 mAb 7-239. While isotype mAb had no inhibitory effect, pre-treatment with 7-239 blocked SARS-CoV-2 uptake (**Figure 3E**), and this Siglec-1 dependency was higher for IFN-α activated APCs. Once Siglec-1 binds HIV-1 or Ebola viruses, this receptor polarizes and engulfs viral particles within VCCs continuous with the plasma membrane and connected to the extracellular space (Izquierdo-Useros et al., 2011; Yu et al., 2008). To elucidate whether the Siglec-1 receptor also recruits SARS-CoV-2 to these compartments, we investigated viral uptake by confocal microscopy. LPS-activated DCs exposed to SARS-CoV-2 showed a Siglec-1 positive VCC containing viral particles attached to the membrane of the compartment (**Figure 3F** and **Supplementary Movie 1**). In addition, using super resolution microscopy of SARS-CoV-2 viral particles stained for the Spike using an ACE2-mFc fusion protein, we confirmed that GM1, one of the sialyllactose-containing gangliosides interacting with Siglec-1 (Izquierdo-Useros et al., 2012a), was detected on 74% of viruses binding the ACE2-mFc fusion protein (**Figure 3G**). Thus, the complementary approaches of Siglec-1 *de novo* expression on Raji cells, combined with the blocking effect of specific mAbs on primary myeloid cells, along with the detection of Siglec-1 interacting GM1 ligands on SARS-CoV-2 particles, supports that Siglec-1 is a central molecule mediating SARS-CoV-2 uptake via sialic acid recognition in myeloid cells.

### 4. Siglec-1 facilitates SARS-CoV-2 trans-infection to target cells

Siglec-1 has a dual role enhancing infectivity of various viruses, either facilitating fusion on APCs, as is the case for Ebola virus, or mediating transmission to other target cells, as is the case for retroviruses. This later mechanism is relevant when APCs are not directly susceptible to infection, as it has been reported for HIV-1 on DCs (Laguette et al., 2011). The inefficient support of SARS-CoV-2 replication on APCs (**Figure 1**) would suggest that Siglec-1 could mediate SARS-CoV-2 transmission to target cells in the context of coronavirus infection. Therefore, we next assessed the relevance of *trans*-infection for SARS-CoV-2 bound via Siglec-1. We pulsed distinct Raji cells with equal amounts of a SARS-CoV-2 Spike pseudotyped HIV-1 construct lacking the lentiviral glycoprotein, but containing a luciferase reporter gene. Pulsed Raji cells were extensively washed and co-cultured with different target cell lines expressing either ACE2 alone or ACE2 and TMPRSS2, a protease required for priming SARS-CoV-2 Spike (Hoffmann et al., 2020). While Raji Siglec-1 cells effectively transferred SARS-CoV-2 to both cellular targets, Raji cells transfected with DC-SIGN marginally transmitted the virus, and cells devoid of any of these two lectins did not transfer SARS-CoV-2 (**Figure 4A**). Moreover, *trans*-infection relied on Siglec-1 uptake of SARS-CoV-2 via recognition of sialylated ligands, as the Siglec-1 mutant R116A did not *trans*-infect SARS-CoV-2 (**Figure 4A**). No direct infection of Raji cells was detected, as seen when assessing pulsed cells in the absence of ACE2-expressing cellular targets. Experiments using wild-type virus confirmed these observations. Raji cells exposed to SARS-CoV-2 or left untreated for 4h were washed and co-cultured with Vero E6 cells at the indicated ratios for 48 h (**Figure 4B**). While Raji Siglec-1 induced higher cytopathic effect on Vero E6 cells, this effect was blocked by Remdesivir treatment and not observed for those co-cultures containing Raji Siglec-1 mutant R116A (**Figure 4B**).

**Figure 4.**
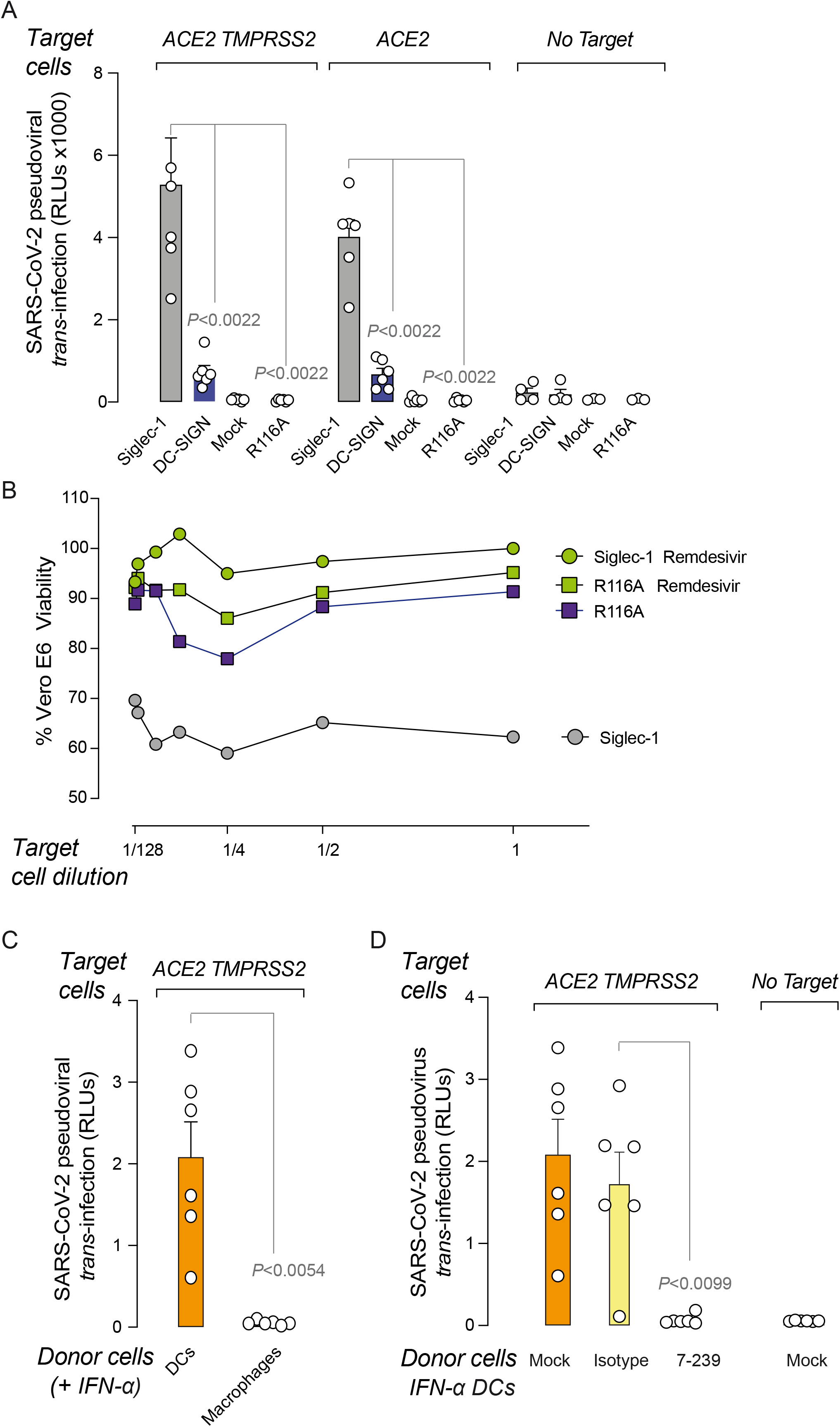
SARS-CoV-2 trapped via Siglec-1 on donor APCs mediates viral *trans*-infection of target cells. **A**. Transmission of SARS-CoV-2 pseudoviruses from distinct Raji cells to the indicated target HEK-293T cells expressing ACE2 or ACE2 and TMPRSS2. Raji cells were pulsed with an HIV-1 luciferase reporter virus lacking the envelope glycoprotein pseudotyped with SARS-CoV-2 Spike for 2 h. Cells were washed and co-cultured with target cells for 48 h. Infection of target cells was determined by induced luciferase activity in relative light units (RLUs). No viral fusion was detected on Raji cells cultured alone. Data show mean values and SEMs from two experiments including three replicates. Statistical differences were assessed with a Mann Whitney test. **B**. Transmission of SARS-CoV-2 by different Raji cells in the presence or absence of 10uM of Remdesivir. Raji cells were exposed to SARS-CoV-2 or left untreated for 4h, washed and cultured with Vero E6 cells at the indicated ratios for 48 h. Percentage of viral induced cytopathic effect was measured by luminometry, and calculated setting co-cultures without viruses at 100% of viability. Data show mean values from one experiment. **C**. Transmission of SARS-CoV-2 pseudoviruses captured for 4 h from DCs and macrophages to HEK-293T cells expressing ACE2 and TMPRSS2. Data show mean values and SEMs from two experiments including cells from six donors. Statistical differences were assessed with a paired t test. **D**. Transmission of SARS-CoV-2 pseudoviruses from IFN-α activated DCs to HEK 293T cells expressing ACE2 and TMPRSS2. Cells were pre-incubated with the indicated mAbs and exposed to SARS-CoV-2 before assessing trans-infection. No viral fusion was detected on DCs not co-cultured with SARS-CoV-2 target cells. Data show mean values and SEMs from two experiments including cells from six donors. Statistical differences were assessed with a paired t test.

We also tested primary monocyte-derived myeloid cells activated with IFNα to up-regulate Siglec-1 expression, and found that DCs where much more efficient at transmitting SARS-CoV-2 when compared to macrophages (**Figure 4C**). We used the α-Siglec-1 mAb 7-239 to explore whether SARS-CoV-2 *trans*-infection could be blocked with this antibody. While isotype mAb had no inhibitory effect, pre-treatment with 7-239 blocked SARS-CoV-2 *trans*-infection to target cells mediated by IFNα-activated DCs (**Figure 4D**). No direct infection of DCs was detected, as seen when assessing pulsed cells in the absence of ACE2-expressing targets. Thus, SARS-CoV-2 retention via Siglec-1 allows *trans*-infection of cells expressing ACE2 and TMPRSS2 receptors, especially by DCs.

### 5. SARS-CoV-2 is detected on APCs expressing Siglec-1 that release inflammatory cytokines in pulmonary tissues of non-human primates

We next interrogated single cell RNA sequencing data (Speranza et al., 2021) previously collected from pulmonary tissues of African Green Monkeys (AGMs) to corroborate our prior *in vitro* observations (**Supplementary Figure 3**). We focused on myeloid APCs and analyzed which of these populations expressed *SIGLEC1* mRNA in the pulmonary tissues of AGMs infected with SARS-CoV-2 over time (**Figure 5A**). We found that tissue-resident alveolar macrophages, DCs, monocyte-derived macrophages or interstial macrophages, monocytes and pDCs all upregulated *SIGLEC1* transcript expression 3 days post-infection (dpi), and that expression was still detectable at 10 dpi, mostly on both types of macrophages (**Figure 5A**). Viral RNA was highly detected at 3 dpi, but decreased at 10 dpi (**Figure 5B**), consistent with the resolution of SARS-CoV-2 infection in these NHP models with mild COVID-19. When we compared the expression of *SIGLEC1* on cells with or without detectable viral RNA, we found that all cells induced *SIGLEC1* transcription upon SARS-CoV-2 infection, although this was significantly more abundant only on alveolar macrophages or interstitial macrophages containing viral RNA, followed by monocytes and DCs, where differences did not reach statistical significance (**Figure 5C**). Indeed, a significant positive correlation between SARS-CoV-2 and *SIGLEC1* transcripts was detected for all APCs but pDCs. Moreover, immunohistochemistry analysis of the pulmonary tissue of a SARS-CoV-2 infected rhesus macaque from a previous study (Hoang et al., 2021) collected at 10 dpi, confirmed at the protein level the co-expression of Siglec-1 and SARS-CoV-2 nucleocapsid in cells with a typical myeloid cell morphology (**Figure 5D**).

**Figure 5.**
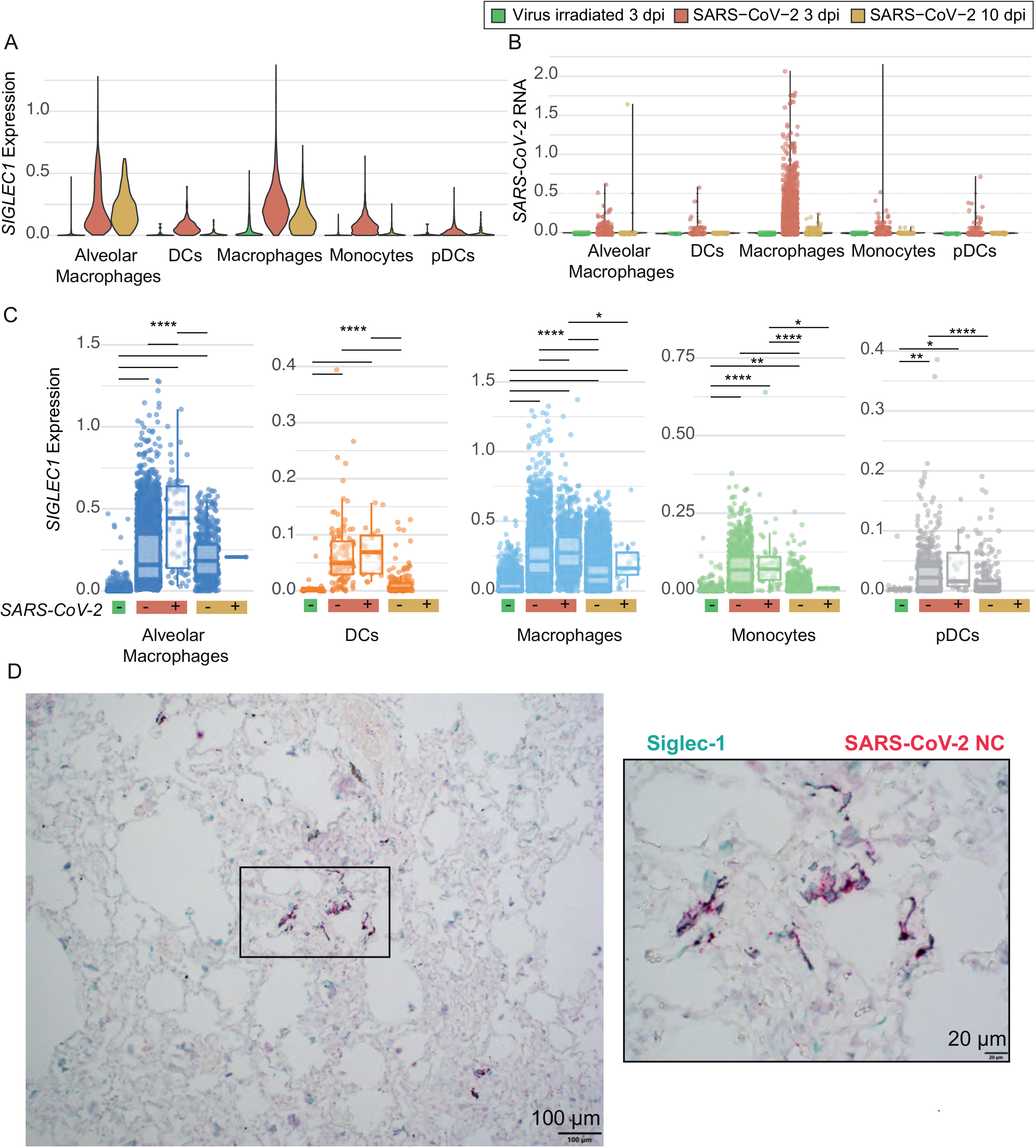
SARS-CoV-2 is detected on APCs expressing Siglec-1 in pulmonary tissues of NHP. **A**. Single cell RNA sequencing (scRNAseq) of *SIGLEC-1* expression on APCs of African Green Monkeys sacrificed after 3 dpi with irradiated virus, or sacrificed after 3 or 10 dpi with SARS-CoV-2. Data from 12 animals. **B**. scRNAseq of SARS-CoV-2 expression as in A, shown as a gene signature encompassing all viral RNA. Data from 12 animals. **C.** SIGLEC-1 expression on APCs containing or not SARS-CoV-2 RNA at different timepoints. Boxplots show median; lower and upper hinges correspond to the first and third quartiles and whiskers extend 1.5 times the interquartile range. Tukey post-hoc test was carried out to test for significance for all pairwise comparisons. Significance levels: ns *P* > 0.05; * *P* ≤ 0.05; ** *P* ≤0.01; *** *P* ≤ 0.001; **** *P* ≤ 0.0001. **D**. Immunohistochemistry of a pulmonary tissue from an infected Rhesus Macaque co-stained with Siglec-1 and SARS-CoV-2 nucleocapsid antibodies. Right panel zooms into the highlighted box.

Finally, we addressed which APCs better sensed SARS-CoV-2 to produce cytokines and IFN-related molecules during the course of the infection in AGMs with mild COVID-19 disease progression (**Figure 6A**). *IL-6* and *IFN-*γ transcripts were significantly triggered by viral presence at 3 dpi in alveolar macrophages or interstitial macrophages and to lesser extend in monocytes and pDCs, but never in DCs (**Figure 6A**). For *IP-10* (CXCL10), however, the induction was also detected for DCs (**Figure 6A**), as we had previously observed by Luminex immunocapture in human cells. For *MCP-1* the increase was evident throughout the course of infection for all APCs, while *IFN-2*α was mostly detected at 3 dpi in interstitial macrophages, monocytes and pDCs not expressing viral RNA (**Figure 6A**). Same trends were observed for all aforementioned molecules in APCs expressing *SIGLEC1* transcripts (**Figure 6B**). When we interrogated GO processes differentially regulated on macrophages containing SARS-CoV-2 RNA, we found that these cells actively modified several processes related to negative regulation of interferon production or regulation of cytokine production involved in immune responses, along with processes involved in antiviral immunity and antigen presentation (**Supplementary Figure 4A**, **Supplementary Table 1**). On DCs, however, processes were related to negative regulation of interferon production, antiviral immunity, and chemotaxis, but not linked to cytokine production (**Supplementary Figure 4B**, **Supplementary Table 2**). Overall, these data indicate that APCs expressing Siglec-1 are found in pulmonary tissues of SARS-CoV-2 infected NHP containing viruses that can trigger cytokine release, which is mostly driven by macrophages.

**Figure 6.**
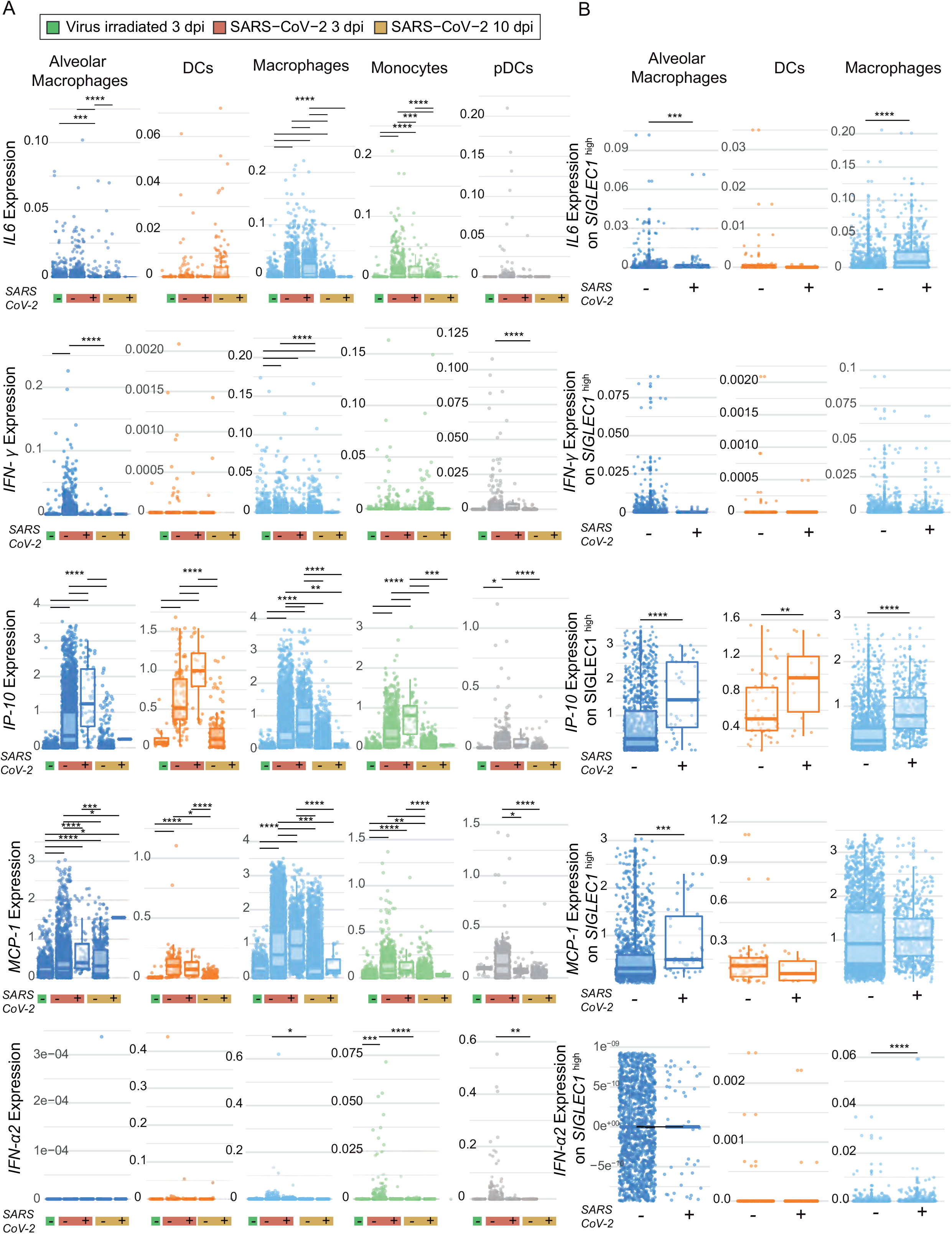
SARS-CoV-2 infection triggers cytokine release mostly in macrophages in pulmonary tissues of NHP. **A**. scRNAseq analysis of *IL-6*, *INF-*γ, *IP-10*, *MCP-1* and *IFN-2*α expression on APCs of African green monkeys sacrificed after 3 dpi with irradiated virus, or sacrificed after 3 or 10 dpi with SARS-CoV-2. Data from 12 animals. Boxplots show median, lower and upper hinges correspond to the first and third quartiles and whiskers extend 1.5 times the interquartile range. **B.***IL-6*, *INF-*γ, *IP-10*, *MCP-1* and *IFN-2*α expression on APCs with high *SIGLEC-1* levels comparing cells with or without SARS-CoV-2 RNA. Data obtained from 12 animals. Boxplots show median, lower and upper hinges correspond to the first and third quartiles and whiskers extend 1.5 times the interquartile range. Tukey post-hoc test was carried out to test for significance for all pairwise comparisons. Significance levels: ns *P* > 0.05; * *P* ≤ 0.05; ** *P* ≤0.01; *** *P* ≤ 0.001; **** *P* ≤ 0.0001.

## DISCUSSION

Here we identified a role for Siglec-1 in SARS-CoV-2 recognition. This IFN-inducible receptor is increased on activated human myeloid cells, and is also detected bearing viruses on APCs of pulmonary tissues of SARS-CoV-2-infected NHP. The association between Siglec-1 expressing APCs and SARS-CoV-2 was also found in peripheral blood monocytes and in the pulmonary lavages of SARS-CoV-2 infected individuals (Bedin et al., 2020; Lempp et al., 2021). In the present study, we define how this interaction takes place via recognition of syalilated gangliosides detected on the viral membrane of SARS-CoV-2 particles. The capacity of Siglec-1 to bind different SARS-CoV-2 variants is consistent with a Spike-independent mode of viral recognition.

Myeloid APCs trapped and engulfed large amounts of SARS-CoV-2 via Siglec-1. Viruses were retained within Sigelc-1 positive VCCs that resemble to those previously described for HIV-1 or Ebola viruses on activated myeloid cells (Izquierdo-Useros et al., 2011; Perez-Zsolt et al., 2019; Yu et al., 2008). Consistent with this *in vitro* findings, Siglec-1 expressing APCs were found in the lungs of SARS-CoV-2 infected NHP. Yet, both activated and steady state APCs were able to continuously process captured viruses in the absence of detectable viral replication. Macrophages secreted higher amounts of cytokines upon viral capture as compared to DCs, indicating that the hyperactivation of these APCs could drive the exacerbated inflammatory cytokine storm-like condition associated with ARDS. These cytokines include IL-6 and TNF-α, which are significant predictors of disease severity and death (Valle, 2020), and MIP1-α, which is elevated in the plasma of intensive care unit patients (Huang et al., 2020). However, in the non-severe SARS-CoV-2 infection NHP model, we failed to detect *TNF-*α nor *MIP1-*α transcripts at 3 dpi, but *IL-6* expression followed the same pattern as that detected *in vitro* on human cells, and was mostly restricted to macrophages. IP-10, an IFN-stimulated gene which is normally up-regulated along Siglec-1 (Rose et al., 2013), is also increased in patients with unfavorable COVID-19 outcomes (Yale IMPACT research team et al., 2020). IP-10 was found upregulated at protein level in human APCs and at transcriptional level in single cell analysis of NHP tissues, both in macrophages and DCs. Finally, and despite the antiviral role of interferons, such as IFN2-α detected both in human APCs and NHP single cell analyses, it is important to highlight that continuous and exacerbated IFN-responses are also linked to severe COVID-19 (Sette and Crotty, 2021; Yale IMPACT Team et al., 2020). On its turn, IFN responses increase Siglec-1 expression, and viral persistence in anatomical reservoirs.

By blocking Siglec-1 function we disrupted SARS-CoV-2 virus binding and uptake, inhibiting viral transmission to target cells. This process, already reported for HIV-1 and CD4^+^ T lymphocytes (Izquierdo-Useros et al., 2012b; Puryear et al., 2013), would involve pulmonary ciliated cells and pneumocytes as the most prominent target cells for SARS-CoV-2 infection. However, here we found that DCs displayed higher *trans*-infection capacity than macrophages. Activated antigen degradation and presentation pathways were more potently induced by SARS-CoV-2 in macrophages than in DCs at transcriptional level in the NHP model, aiding to explain this observation. Siglec-1 is a dispensable host factor which is absent in rare null homozygous individuals (Martinez-Picado et al., 2016) that interacts with viral gangliosides (Izquierdo-Useros et al., 2012a, 2012b; Puryear et al., 2012, 2013). Thus, targeting this receptor can mitigate potential viral evasion mechanisms observed for neutralizing mAbs directed against viral glycoproteins of different viruses. Further work should address the effect of antibodies targeting Siglec-1 on SARS-CoV-2 dissemination and systemic spread of distinct variants in relevant animal models. It is critical to also measure the possible implication of this receptor in the induction of antiviral immune responses against SARS-CoV-2. Finally, these studies could pave the way to understand if Siglec-1 modulation with mAbs could decrease cytokine storm implicated in ARDS associated to severe COVID-19, as recently suggested for other lectin receptors (Lu et al., 2021). Since the capture of very distant viruses such as HIV-1, Ebola virus and now SARS-CoV-2 is regulated by Siglec-1, strategies targeting this receptor may offer new broad-spectrum inhibitors for enveloped viruses, which may be even effective for future outbreaks.

## MATERIAL AND METHODS

### Ethics statement

The institutional review board on biomedical research from Hospital Germans Trias i Pujol (HUGTiP) approved this study. The biologic biosafety committee of the Research Institute Germans Trias i Pujol approved the execution of SARS-CoV-2 experiments at the BSL3 laboratory of the Center of Bioimaging and comparative imaging (CMCIB).

### Cell cultures

Vero E6 cells (ATCC CRL-1586) were cultured in Dulbecco’s modified Eagle medium, (DMEM; Lonza) supplemented with 5% fetal bovine serum (Invitrogen), 100 U/mL penicillin, 100 μg/mL streptomycin, and 2 mM glutamine (all ThermoFisher Scientific). HEK-293T (ATCC repository) were maintained in DMEM with 10% fetal bovine serum, 100 IU/mL penicillin and 100 μg/mL streptomycin (all from Invitrogen). Raji B lymphocyte and Raji DC-SIGN cell lines (kindly provided by Y. Van Kooyke) were maintained in Roswell Park Memorial Institute medium (RPMI; Invitrogen) or RPMI plus 1 mg/mL geneticin (Invitrogen). Generation and maintenance of Raji Siglec-1, Raji R116A, Raji Siglec-5 and Raji Siglec-7 has been described elsewhere (Perez-Zsolt et al., 2019). HEK-293T overexpressing the human ACE2 were kindly provided by Integral Molecular Company and maintained in DMEM with 1 μg/mL of puromycin (Invitrogen). TMPRSS2 human plasmid (Origene) was transfected using X-tremeGENE HP Transfection Reagent (Merck) on HEK-293T overexpressing the human ACE2 and maintained in the previously described media containing 1 mg/mL of geneticin (Invitrogen) to obtain TMPRSS2/ACE2 HEK-293T cells. All media contained 10% fetal bovine serum, 100 IU/mL penicillin and 100 μg/mL streptomycin(all from Invitrogen).

### Primary Cell Cultures

Peripheral blood mononuclear cells were obtained with a Ficoll-Hypaque gradient (Alere Technologies AS) from blood donors and monocyte populations (>90% CD14^+^) were isolated with CD14-negative selection magnetic beads (Miltenyi Biotec). Macrophages were obtained culturing these cells in the presence of 100 μg/mL of macrophage colony-stimulating factor (M-CSF) for seven days and replacing media and cytokines every 2 days. DCs were obtained culturing these cells in the presence of both 1,000 IU/mL of granulocyte-macrophage colony-stimulating factor (GM-CSF) and interleukin-4 (IL-4; both from R&D) for seven days and replacing media and cytokines every 2 days. Activated cells were differentiated by culturing myeloid cells at day five for two more days in the presence of 1,000 IU/mL of interferon-alfa (IFNα; Sigma-Aldrich) or 100 ng/mL of lipopolysaccharide (LPS, Sigma-Aldrich).

### Virus Isolation, Titration and Sequencing

Unless otherwise specified, SARS-CoV-2 used was the virus isolated in March 2020 from a nasopharyngeal swab as described in (Rodon, 2021). The virus was propagated for two passages and a virus stock was prepared collecting the supernatant from Vero E6. Genomic sequence was deposited at GISAID repository (http://gisaid.org) with accession ID EPI_ISL_510689. Compared to the Wuhan/Hu-1/2019 strain, this isolate has the following point mutations: 376 D614G (Spike), R682L (Spike), and C16X (NSP13). The SARS-CoV-2 B.1.1.7 variant (originally isolated from the UK), the B.1.1.248.2 variant (originally isolated from Brazil) and the B.1.351 variant (originally isolated from South Africa) were identified during routine sequencing of a clinical nasopharyngeal swabs in Spain during January-February 2021 and subsequently isolated on Vero E6 cells. These sequences are deposited at GISAID database with accession numbers EPI_ISL_1663567; EPI_ISL_1831696 and EPI_ISL_1663571 for B.1.17, B.1.1.248.2 and B.1.351, respectively. Genomic sequencing was performed from viral supernatant by using standard ARTIC v3 based protocol followed by Illumina sequencing (dx.doi.org/10.17504/protocols.io.bhjgj4jw). Raw data analysis was performed by viralrecon pipeline (https://github.com/nf-core/viralrecon) while consensus sequence was called using samtools/ivar at the 75% frequency threshold.

### Pseudovirus production

HIV-1 reporter pseudoviruses expressing SARS-CoV-2 Spike protein and luciferase were generated using two plasmids. pNL4-3.Luc.R-.E-was obtained from the NIH AIDS repository. SARS-CoV-2.SctΔ19 was generated (Geneart) from the full protein sequence of SARS-CoV-2 spike with a deletion of the last 19 amino acids in the C-terminal, human-codon optimized and inserted into pcDNA3.4-TOPO (Ou et al., 2020). Spike plasmid was transfected with X-tremeGENE HP Transfection Reagent (Merck) into HEK-293T cells, and 24 h post-transfection, cells were transfected with pNL4-3.Luc.R-.E-. VSV-G plasmid (kindly provided by A. Cimarelli) was used to equally pseudotype pseudoviruses. Supernatants were harvested 48 h later, filtered with 0.45 μM (Millex Millipore) and stored at −80°C until use. Viruses were titrated in HEK-293T overexpressing the human ACE2.

### Pseudoviral fusion assay

Macrophages or DCs activated or not with IFNα as previously described along with HEK-293T ACE2 cells were exposed to equivalent MOI of VSVg or SARS-CoV-2 spike pseudotyped lentiviruses. To block ACE2 dependent viral fusion, some wells had 20 μg/mL of human ACE2-murine Fc fusion protein. Two days post-infection, cells were lysed with the Glo Luciferase system (Promega). Luminescence was measured with an EnSight Multimode Plate Reader (Perkin Elmer).

### Construction of a human-ACE2 murine-Fc-fusion protein (ACE2-mFc)

Expression vector was generated with the Geneart service (Thermofisher Scientific). Coding sequence included the first 615 amino acids from the Human ACE2 sequence, with H345A and H505A mutations to inactivate the catalytical sites, followed by the constant region of the heavy chain of the murine IgG1. For protein production, Expi293F cells (Thermofisher Scientific) were transfected with ACE2-mFc vector at a density of 2.5 × 10^6^ cells/mL using Expifectamine (Thermofisher Scientific). Enhancers 1 and 2 (Thermofisher Scientific) were added to the culture 18 h post-transfection. Cells were incubated for 5 days and supernatants were harvested and passed through a 0.22 μm PVDF filter. For purification, supernatants were loaded into a 5 mL SepFast Ø11mm (Quimigen) packed with CaptureSelect™ IgG-Fc (ms) affinity resin (Thermo Scientific) connected to an Äkta Start Chromatograph (Cytiva). The column was washed with 5 column volumes (CV) of PBS and ACE2-mFc was eluted with 2 CV of 0.1M Glycine at pH=3.5. The sample was concentrated with a 30kDa Amicon Centrifugal Concentrator at 3000 × g. ACE2-mFc concentration was determined by sandwich ELISA using a goat anti-mouse IgG Fc (Jackson Immunoresearch, 115-006-071) for capture, a horseradish peroxidase (HRP) labeled F(ab)2 Goat anti-mouse IgG Fc (Jackson immunoresearch, 115-036-071) as secondary antibody, A purified mouse IgG (D50, NIH AIDS Reagent Program) as standard and o-phenylenediamine dihydrochloride (Sigma-Aldrich, #P8787-100TAB) as substrate. Light absorbance was measured at 492/620 nm on EnSight Multimode Plate Reader (Perkin Elmer). ACE2-mFc inhibitory capacity was tested in a SARS-CoV-2 neutralization assay as described previously (Trinité et al., 2021).

### SARS-CoV-2 Uptake and Degradation Assays

Uptake experiments with SARS-CoV-2 were performed pulsing 0.5 × 10^6^ Raji or 1 × 10^6^ myeloid cells at a rate of 70 ng of Nucleocapsid at 37°C for the indicated timepoints. For blockade, cells were pre-incubated for 15 min at RT with 10 μg/mL of mAbs α-Siglec-1 7–239, or IgG1 isotype control (BD Biosciences), or left untreated before viral exposure. After extensive washing, cells were lysed at a constant concentration of 1×10^6^ cells/mL, centrifuged to remove cellular debris and assayed with a SARS-CoV-2 Nucleocapsid protein (NP) High-sensitivity Quantitative ELISA (ImmunoDiagnostics). For degradation experiments, myeloid cells were exposed to SARS-CoV-2 for 4h, extensively washed, and left in culture for the indicated timepoints until cell associated viral content and viral release to the supernatant were measured with the indicated ELISA kit.

### Electron microscopy of myeloid cells

10 × 10^6^ myeloid cells (macrophages or DCs activated with LPS) were exposed to SARS-CoV-2 with an MOI of 1 for 24h, fixed with paraformaldehyde (PFA) at 4% (Biotium) and glutaraldehyde 1% (Sigma Aldrich/Merck) for one hour at room temperature (RT) and processed for embedding in resin, ultramicrotomy and transmission electron microscopy as previously described (Tenorio et al., 2018). Briefly, after fixation the samples were washed three times with PBS and cells were gently scraped with a rubber policeman. Cell pellets were postfixed with 1% osmium tetroxide + 0.8% potassium ferrocyanide in water for 1h on ice. The samples were dehydrated on ice with a gradual series of acetone and infiltrated at RT with epoxy resin. After heat polymerization the samples were sectioned with a UC6 microtome with a nominal feed of 70 nm. Sections were collected on 300 mesh bare copper grids and contrasted with 4% aqueous uranyl acetate, followed by Reynold’s lead citrate. Images were taken either using a Jeol 1011 run at 100 kV equipped with a Gatan ES1000W camera or a Jeol 1400 run at 80kV with a Gatan Oneview camera.

### Cytokine profiling

Cytokines were measured by Luminex xMAP technology and analyzed with xPONENT 3.1 software (Luminex Corporation) using the MCYTOMAG-70 kit, according to the protocol of the manufacturer, with minor modifications. Briefly, after staining the desired cytokines, an overnight incubation was performed in a rocking shaker at 4°C using 2% PFA to fully inactivate remaining SARS-CoV-2 particles, a fixation that does not alter quantification (Dowall et al., 2009). Before plate acquisition PFA was washed and changed for sheath fluid.

### Activation of myeloid cells

Myeloid cells were left untreated activated for 48h with SARS-CoV-2 at a MOI of 0.1 and compared to cells treated with IFNα or LPS as previously described. Cells were blocked with 1 mg/mL human IgG (Privigen, Behring CSL) and stained with anti-Siglec-1-PE 7-239 mAb (AbD Serotec), anti-DC-SIGN-PE DCN46 mAb, anti-HLA-DR-PerCP L243 mAb, anti-CD86-FITC 2331 (FUN-1) mAb, anti-CD83-FITC HB15e mAb and anti-CD14-PerCP MφP9 mAb (all from BD Biosciences) at 4 °C for 30 min. Mouse IgG1-PE (AbD Serotec) was included as isotype control. Samples were analyzed with FACS Canto (BD Biosciences) using FlowJo software to evaluate collected data.

### Relative mRNA quantification

RNA extraction was performed using MagMAX™ mirVana total RNA isolation Kit, optimized for a KingFisher instrument (ThermoFisher) Reverse transcriptase was performed using the PrimeScript™ RT-PCR Kit (Takara). mRNA relative levels of *SIGLEC1* were measured by two-step quantitative RT-PCR and normalized to GAPDH mRNA expression using the DDCt method. Primers and DNA probes were purchased from Life Technologies TaqMan gene expression assays.

### Confocal Microscopy analyses

LPS-treated DCs were pulsed with SARS-CoV-2 with an MOI of 1 for 4 h at 37 °C. After extensive washing, cells were fixed and permeabilized (Fix & Perm, Invitrogen) and stained with anti-rabbit nucleocapsid pAb (GeneTex) revealed with a Goat pAb Anti-Rabbit IgG Alexa 488 (Abcam) and an anti-Siglec-1 7-239 Alexa 647 mAb (Biolegend). Cells were cytospun into coverslips, covered with DAPI-containing Fluoroshield mounting medium (Sigma-Aldrich) and analyzed with a Dragonfly (Andor) 505 multimodal confocal microscope with GPU driven deconvolution to maximize resolution.

### Super-resolution analysis of SARS-CoV-2

For super-resolution detection of nucleocapsid and Spike proteins and GM1 gangliosides, SARS-CoV-2 particles were adhered to poly-L coated coverslips for 15 min at RT and fixed in 4 % PFA/PBS for 30 min. Fixed and inactivated virus samples were permeabilized and blocked using 0.1 % saponin /0.5 % BSA/PBS. Virus particles were immunostained with rabbit anti-SARS-CoV-2 N protein (Sino Biological) or rabbit anti-GM1 Ab (Abcam) followed by anti-rabbit Abberior STAR RED (Abberior GmbH) Fab fragments. SARS-CoV-2 Spike protein was detected with an ACE2-mAb Fc recombinant protein and anti-mouse Abberior STAR 580 (Abberior GmbH) Fab fragments. SARS-CoV-2 Spike protein was detected with an ACE2-mAb Fc recombinant protein and anti-mouse Abberior STAR RED conjugated Fab fragments. Following immunostaining, all samples were overlaid with SlowFade Diamond mounting medium (ThermoFisher Scientific, USA) and imaged using STED microscopy. Super-resolution analysis of SARS-CoV-2 virus particles was performed using Leica SP8 STED 3X microscope (Mannheim, Germany) equipped with a 100×/1.4 NA oil immersion STED objective. STED images of N protein, GM1 (Abberior STAR RED) and S-ACE2 protein complexes (Abberior STAR 580) signals were acquired sequentially for each channel using 637 nm and 587 nm lines from the white light laser. Abberior STAR RED and Abberior STAR 580 signal was depleted with a donut-shaped 775-nm pulsed STED laser. STED depletion conditions were tuned to achieve 40 nm lateral resolution (full-width-at-half-maximum, FWHM) as estimated from fluorescent bead and single fluorescent antibody molecule measurements. STED images were acquired with following parameters: pinhole size: 1.03 Airy; dwell time: 2 μs/pixel and XY pixel size: 20 nm. Acquired STED images were thresholded and filtered using Gaussian filter (Sigma (Radius) = 0.75) using Fiji (ImageJ distribution) software.

### Trans-infection Assay

HEK-293T overexpressing the human ACE2, or both ACE2 and TMPRSS2 were used to test if SARS-CoV-2 pseudotyped viruses can fuse with these receptors via *trans*-infection. A constant pseudoviral titer was used to pulse cells in the presence of the indicated mAbs for 2 h at 37°C. Cells were extensively washed and co-cultured with or without target cells at a ratio 1:1. Two days post-infection, cells were lysed with the Glo Luciferase system (Promega). Luminescence was measured with an EnSight Multimode Plate Reader (Perkin Elmer).

Vero E6 cells were used to test if SARS-CoV-2 can be *trans*-infected. A constant MOI of SARS-CoV-2 from B.1.1.7 variant was used to pulse Raji cells for 4 h at 37°C. Cells were extensively washed and co-cultured with or without target cells at the indicated ratios. One duplicate was co-cultured in the presence of 10 μM of Remdesivir to block viral replication. Cells not exposed to the virus were equally co-cultured with Vero E6 cells and used to set 100% of cell viability for each condition. Two days post-infection, cells were lysed with the Cell Titter Glo viability system (Promega). Luminescence was measured with a Luminoskan (ThermoScientific).

### Data origin, quality control and data processing of single-cell RNA sequencing methods

We used publicly available lung single-cell RNA sequencing data from Chlorocebus aethiops (African green monkey)[10.1126/scitranslmed.abe8146] in which monkeys were inoculated with infectious SARS-CoV-2 or irradiated, inactivated virus and sacrificed at 3dpi and 10dpi. The data originally used by Speranza *et al.* [10.1126/scitranslmed.abe8146] is publicly available in the Gene Expression Omnibus through GEO Series accession no. GSE156755 (www.ncbi.nlm.nih.gov/geo/query/acc.cgi?acc=GSE156755). Data generation protocols and read mapping can be found in the original publication. Raw data was loaded and processed with R 4.0.1 [R Core Team (2021). R: A language and environment for statistical computing. R Foundation for Statistical Computing, Vienna, Austria. URL https://www.R-project.org/] using Seurat V4 [https://doi.org/10.1101/2020.10.12.335331]. Processing and quality control steps were carried out as detailed in the original manuscript, all datasets were integrated using Seurat’s IntegrateData function and we then filtered out low quality cells and doublets by removing cells with abnormally mitochondrial genes (greater than 3 SDs above the median), and cells that were likely doublets were relabeled [ratio of unique features to unique mapped identifier (UMI) per cell < 0.15]. Also, cells containing less than or greater than 3 SDs of UMI compared to the population total were removed to filter for noise. [10.1126/scitranslmed.abe8146]. Principal component analysis was carried out in the integrated space and clustering and UMAP embedding was performed using the top 30 principal components. Data normalization was carried out in the raw count matrix space using Seurat’s function SCTransform.

### Single-cell RNA sequencing downstream analysis and code availability

Cells were annotated using canonical marker genes obtained from the original paper and previously published human lung atlases [https://doi.org/10.1038/s41586-020-2922-4]. A SARS-CoV-2 gene signature looking at all the viral RNAs was obtained for all the cells and those with a positive score, identifying those with at least one viral RNA, were labeled as SARS-CoV-2-positive. Due to the sparsity of the scRNAseq data we used MAGIC [https://doi.org/10.1016/j.cell.2018.05.061] denoising on the normalized count matrix to recover the underlying structure of the data and enhance the signal of lowly expressed genes. To statistically test the levels of *SIGLEC1* at the different necropsy timepoints and between SARS-CoV-2^+^ and SARS-CoV-2^-^ we carried out pairwise Tukey post-hoc tests across all conditions. The same statistical test was carried out when assessing expression-level changes of key cytokines across conditions. Lastly GO enrichment analysis was carried out between populations of interest: SARS-CoV-2^+^ macrophages vs SARS-CoV-2^-^ macrophages and SARS-CoV-2^+^ DCs vs SARS-CoV-2^-^ DCs. To carry out GO enrichment we first computed differentially expressed genes between the populations of interest using Seurat’s function FindMarkers to define our gene set and we used entire gene space as the gene universe set. We used the GOstats packages to obtain the enriched GO terms using the human annotation as the reference. To narrow down the enriched GO terms to relevant ones we filtered significant GO terms with a size between 5 and 100 genes. All analyses were carried out using R4.0.1[R Core Team (2021). R: A language and environment for statistical computing. R Foundation for Statistical Computing, Vienna, Austria. URL https://www.R-project.org/] and data was analyzed using Seurat V4 [https://doi.org/10.1101/2020.10.12.335331]. All data and code can be found in the GitHub repo https://github.com/MarcElosua/SIGLEC1-SARS-CoV-2

### Immunohistochemical staining on sections of SARS-CoV-2

The paraffin-embedded sections of SARS-CoV-2 infected lungs from a previous study with Rhesus macaques (Hoang et al., 2021) were subjected to deparaffinization in xylene, rehydration in graded series of ethanol, and rinsed with double distilled water. Antigen retrieval was performed by immersing sections in DIVA Decloaker (Biocare Medical) at 125°C for 30 seconds in a steam pressure decloaking chamber (Biocare Medical) followed by blocking with SNIPER Reagent (Biocare Medical) for 10 min. The sections were incubated with SARS Nucleocapsid Protein Antibody (Rabbit polyclonal; Novus Biologicals, NB100-56576SS) and rabbit anti-human Anti-Sialoadhesin/CD169 antibody (clone SP216, Abcam, ab183356) for 1 h, followed by a double detection polymer system (Mach 2 Double Stain 2, Biocare Medical). Labeled antibodies were visualized by development of the chromogen (Warp Red and/or Vina Green Chromogen Kits; Biocare Medical). Digital images of lung were captured at 100×, 200× and 400× magnification with an Olympus BX43 microscope equipped with a digital camera (DP27, Olympus) and evaluated using Cellsens Standard digital imaging software 2.3 (Olympus).

### Statistical analyses of non-single cell data

Statistical differences from 100% of viability were assessed with a one-sample t-test. Statistical differences were also assessed with a Mann Whitney *t* test, a Wilcoxon matched paired *t* test and a paired t test. Comparisons were performed with Graph Prism 9.

## SUPPLEMENTARY FIGURES

**Supplementary Figure 1**. Weak activation of myeloid cells in the presence of SARS-CoV-2. Representative staining of the indicated markers of DCs and Macrophages mock treated or exposed to SARS-CoV-2 (MOI=0.01), LPS (100ng/mL) and IFN-α (1000 U/mL) for 48h before cell staining, analyses by FACS. Results from three independent biological replicates and one experiment.

**Supplementary Figure 2.** Representative surface staining of the different lectins expressed on transduced Raji cell lines analyzed by FACS.

**Supplementary Figure 3. A.** Uniform Manifold Approximation and Projection embedding of the cells analyzed by scRNAseq colored by their annotated cell type. **B**. DotPlot showing the marker gene expression for each cell type.

**Supplementary Figure 4**. **A**. Enriched GO terms in macrophages containing SARS-CoV-2 RNA as compared to macrophages lacking viral RNA. Statistical differences were assessed with a hypergeometric test. **B**. Enriched GO terms differentially regulated on DCs as in A.

## SUPPLEMENTARY TABLES

**Supplementary Table 1.** List of GO processes differentially regulated on macrophages containing SARS-CoV-2 RNA as compared to those lacking viral RNA.

**Supplementary Table 2.** List of GO processes differentially regulated on DCs containing SARS-CoV-2 RNA as compared to those lacking viral RNA.

## SUPPLEMENTARY MOVIES

**Supplementary Movie 1**. Deconvolved 3D reconstruction of an LPS-treated DC exposed to SARS-CoV-2 showing a VCC. Green; anti-nucleocapsid pAbs, Red; anti-Siglec-1 mAb, and Blue; Dapi staining the nucleus.

## AUTHOR CONTRIBUTION

Conceived and designed the experiments: B.C, J.M-P, J.V-A, N.I-U.

Performed experiments and provided key reagents: D.P-Z, J.M-B, J.R, M. E, D.R-R, C.R, M.S, M.P., S.G., MPa, J.C, E.B, E.R, I. E, X. M, M. N, J.B, J.S, J.V-A, N.I-U.

Analyzed and interpreted the data: D.P-Z, J.M-B, J.R, M.E., D.R-R, C.R, M.S, M.S, M.P., S.G., J.C, E.B, E.R, I. E, X. M, M. N, R. P, I. B, V. G, J.C, J.B, A.T., H.H., J.S, B.C, J.M-P, J.V-A, N.I-U.

Wrote the paper: D.P-Z, J.M-B, J.R, M.E., D.R-R, J.V-A, N.I-U.

## ACKNOWLEDGEMENTS

We acknowledge J. Pedroza from the CMCIB for his constant help at the BSL3 facility. We thank C. Esteban from the microbiology department for identifying two different variants used in this study. We also thank M. Parera from IrsiCaixa for her outstanding help in sequencing viral variants. We are grateful to M. Fernandez of the IGTP Cytometry platform for his excellent assistance for Luminex acquisition and analysis, and to E. Rebollo from the Advanced Fluorescence Microscopy Unit IBMB-PCB. We would like to thank Advanced Light Microscopy Unit at the Centre for Genomic Regulation (CRG), Barcelona, Spain for the access to Leica STED microscope.

## DATA AVAILABILITY

Data is available from corresponding author upon reasonable request.

