## Supplementary figures and images for "Siglec-1 on dendritic cells mediates SARS-CoV-2 *trans*-infection of target cells while on macrophages triggers proinflammatory responses"

### Supplementary Figure 1

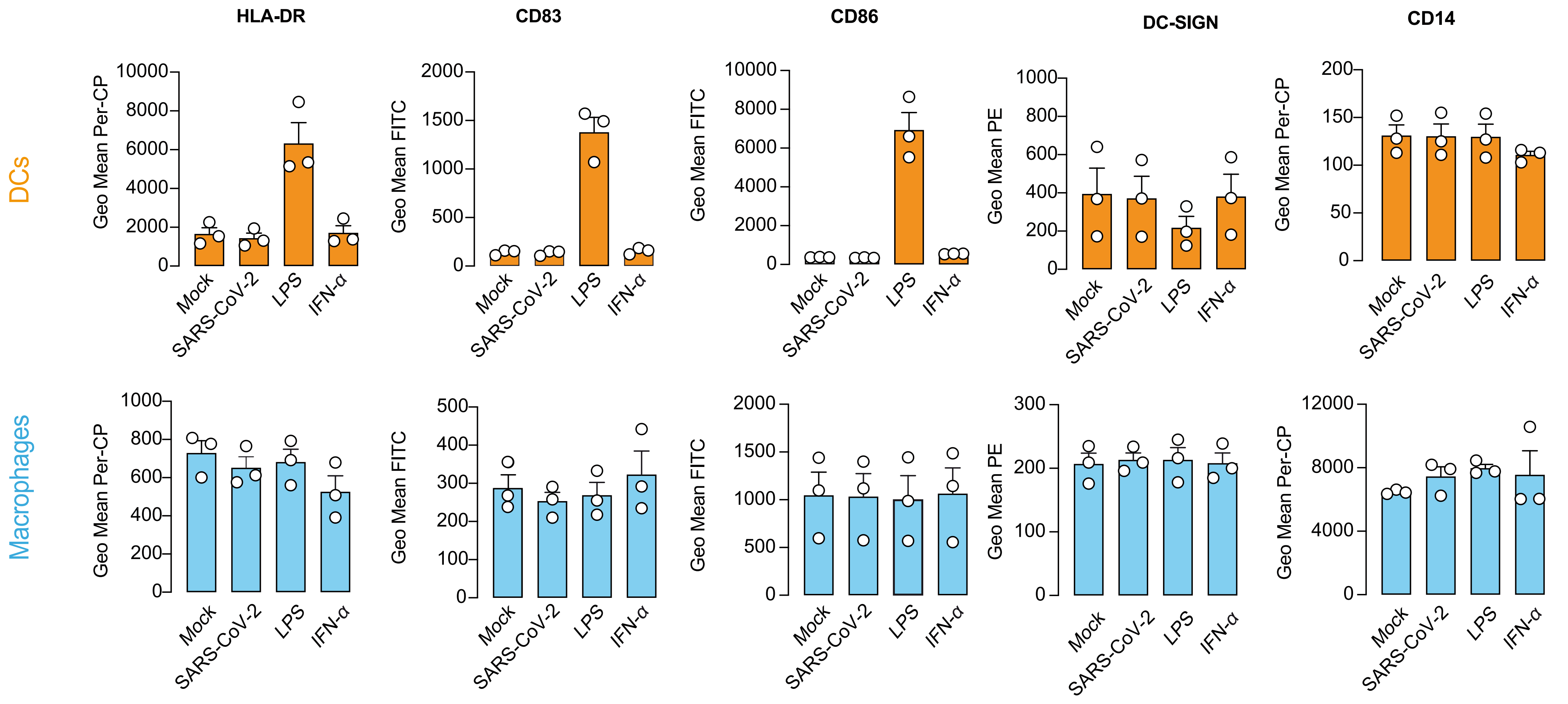


Supplementary Figure 1

### Supplementary Figure 2

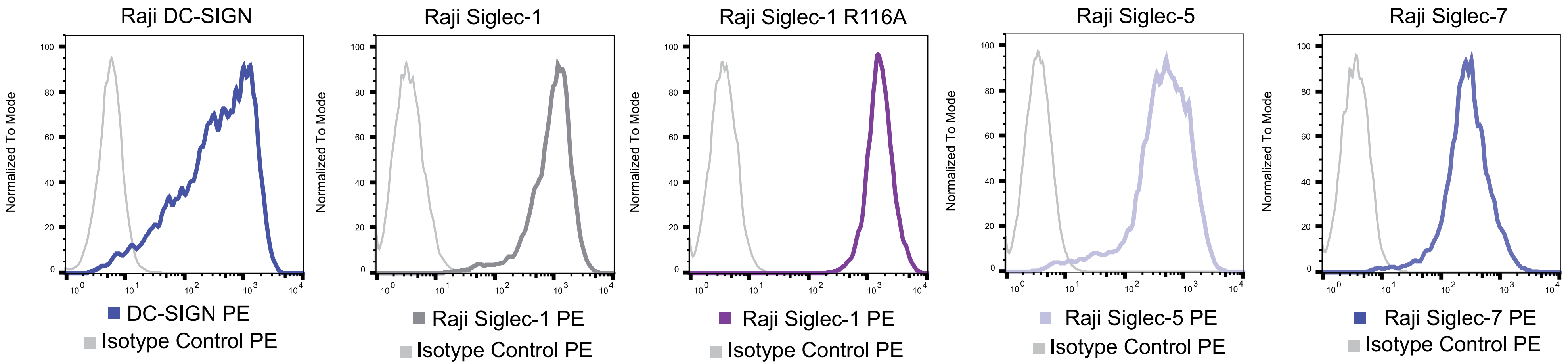


Supplementary Figure 2

### Supplementary Figure 3

A

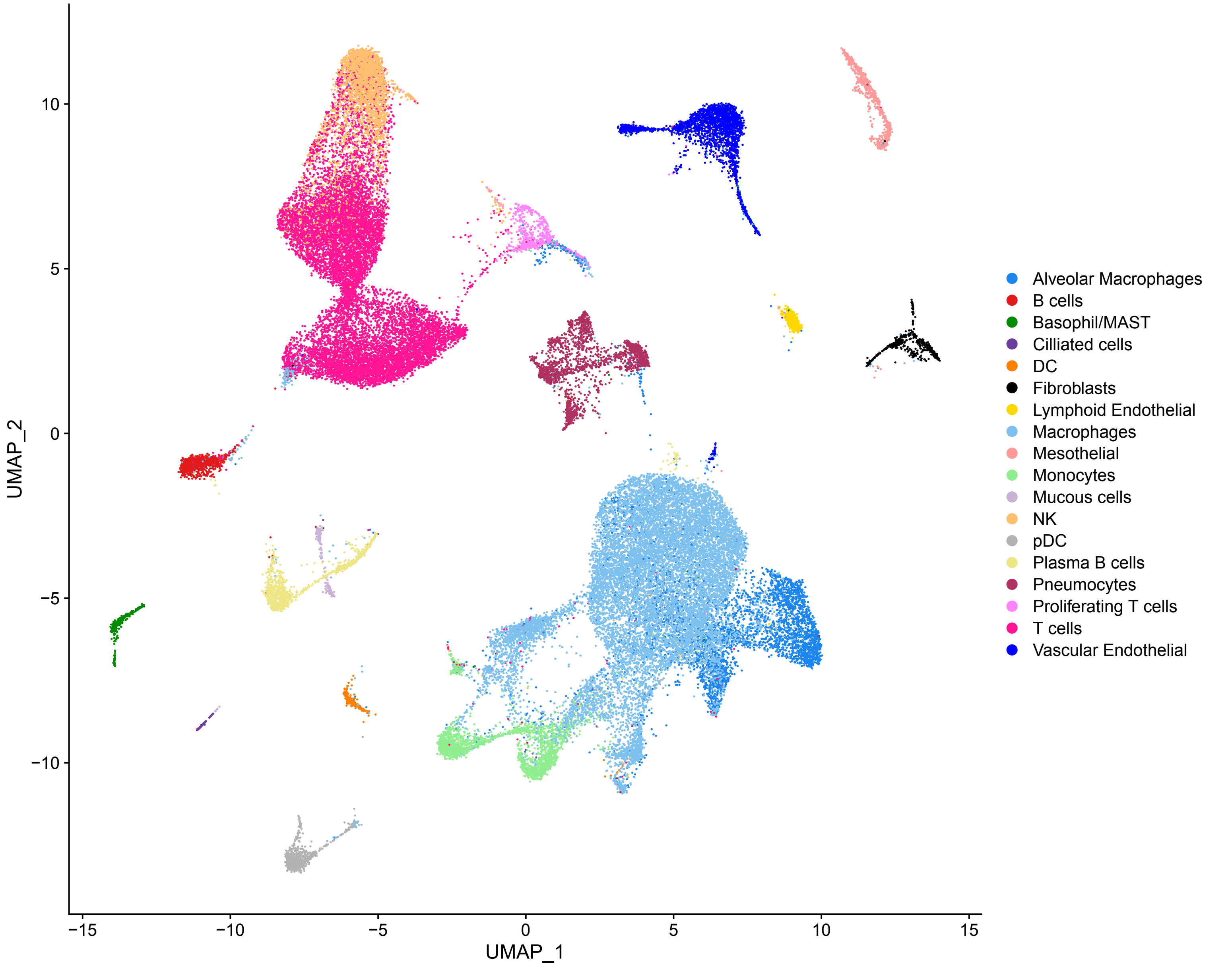


B

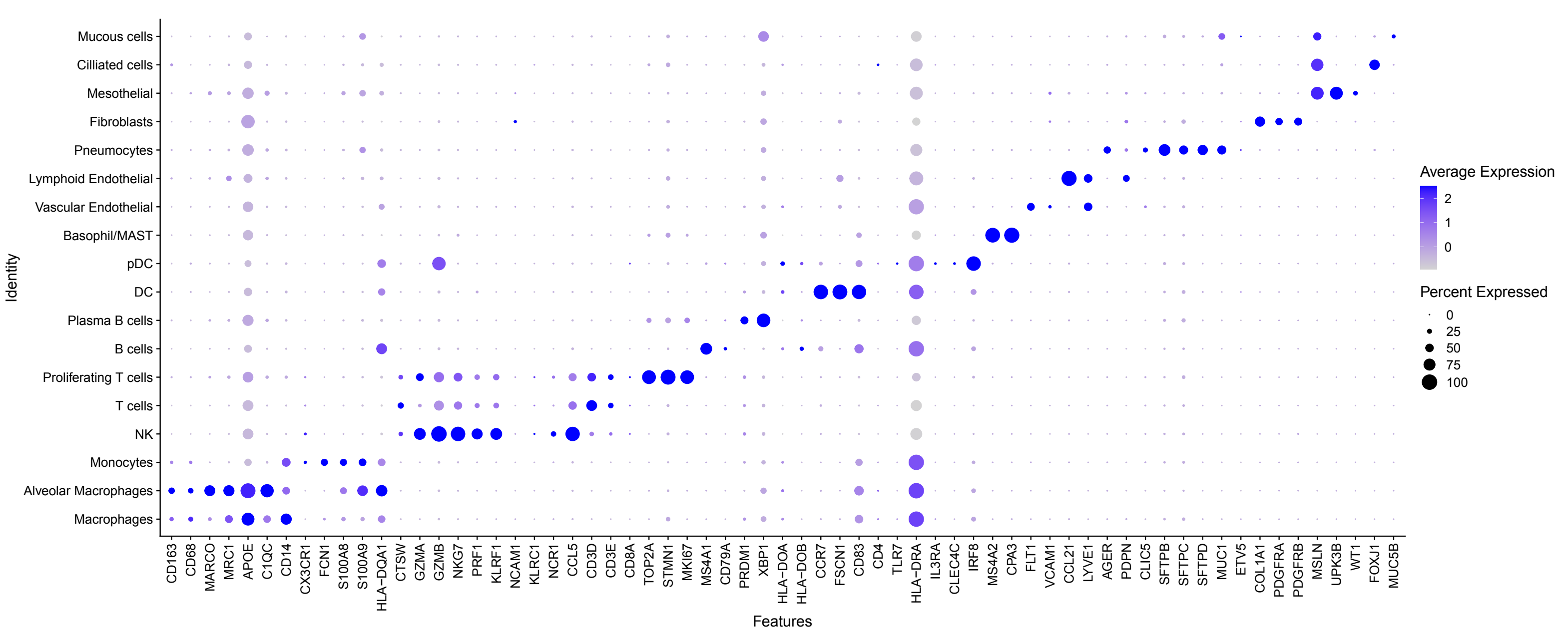


Supplementary Figure 3

### Supplementary Figure 4

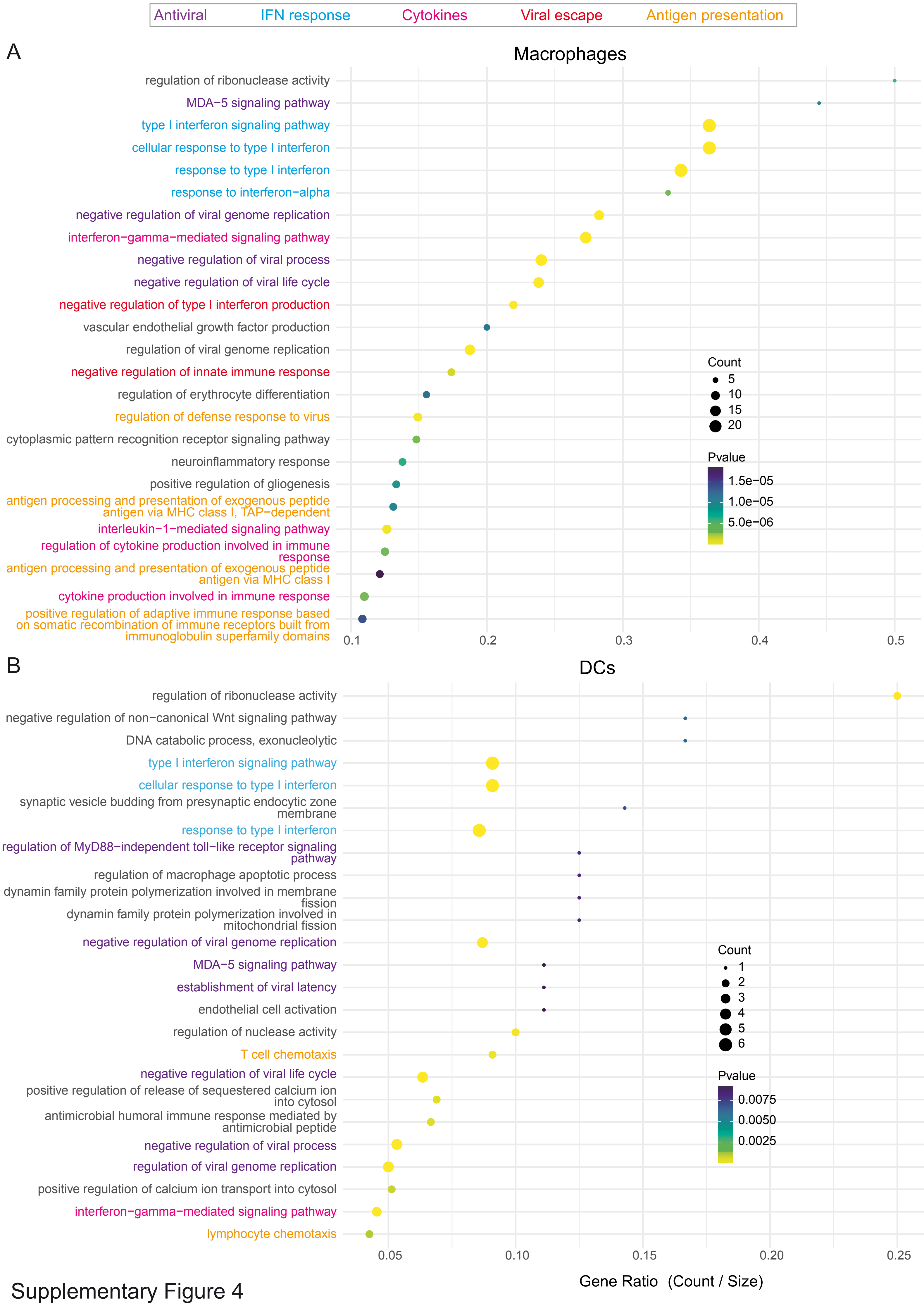


Supplementary Figure 4
